# scGRNom: a computational pipeline of integrative multi-omics analyses for predicting cell-type disease genes and regulatory networks

**DOI:** 10.1101/2020.06.11.147314

**Authors:** Ting Jin, Peter Rehani, Mufang Ying, Jiawei Huang, Shuang Liu, Panagiotis Roussos, Daifeng Wang

## Abstract

Understanding cell-type-specific gene regulatory mechanisms from genetic variants to diseases remains challenging. To address this, we developed an open-source computational pipeline, scGRNom, to predict the cell-type disease genes and regulatory networks from multi-omics data, including cell-type chromatin interactions, epigenomics, and single-cell transcriptomics. With applications to Schizophrenia and Alzheimer’s Disease, our predicted cell-type regulatory networks link transcription factors and enhancers to disease genes for excitatory and inhibitory neurons, microglia, and oligodendrocytes. The enrichments of cell-type disease genes reveal cross-disease and disease-specific functions and pathways. Finally, machine learning analysis found that cell-type disease genes shared by diseases have improved clinical phenotype predictions.

## Background

Recent Genome Wide Association Studies (GWAS) studies have identified a variety of genetic risk variants associated with multiple brain diseases. For example, a recent study found 109 pleiotropic loci significantly associated with at least two brain disorders [1]. Many cross-disease common genetic risk factors have revealed many shared functional consequences in clinical presentations [2]. Recent studies have also revealed shared symptoms at both psychiatric and physical levels between neurodegenerative and neuropsychiatric diseases [3]. For instance, 97% of Alzheimer’s disease patients develop neuropsychiatric symptoms throughout the disease [4]. Besides, additional insights into each disease’s progression and causes have further demonstrated the highly interlinked nature of both disease types [5]. However, our understanding of the molecular mechanisms of genetic variants between diseases remains elusive, particularly at the cell-type levels.

Alzheimer’s disease (AD) and Schizophrenia (SCZ) are representative neurodegenerative and neuropsychiatric diseases, respectively. Both are significantly associated with genetic variants and have complex underlying cellular and molecular mechanisms from genotype to phenotype [6,7]. Notably, AD is physiologically characterized by accumulations of amyloid beta plaques and neurofibrillary tau protein tangles in the brain. Amyloid beta plaques primarily originate from the apolipoprotein E encoding gene APOE and its multiple variants. The APOE gene is a single step in the broader amyloidogenic processing pathway (APP), and additional genes involved in the process contribute to the regulation of amyloid beta production. Much work has identified major genes of interest involved in the APP. However, a distinct need still exists to further explore these disease loci to understand better the interplay between their regulatory elements and eventual amyloid beta creation and accumulation. Similarly, neurofibrillary tau tangles are associated with many genetic loci and require a study of the highly complex molecular mechanisms required to achieve disease pathology. Further, the downstream effects from both amyloid beta and neurofibrillary tangles within and between various cell types add additional complexity toward linking specific regulatory events and elements with clinical pathology [8,9].

Also, SCZ is a neuropsychiatric disorder characterized by disruptions in dopamine, glutamate, and GABA-based receptor signaling pathways. Pathologically, the direct connection is less clear between known molecular abnormalities and observed physical changes through neurological imaging studies. Thus, attempting to understand interactions between activation of known risk genes and higher-level pathway disruptions may help elucidate causes for structural shifts in SCZ patients. At a psychiatric level, the alterations to various cortical structures create multiple forms of symptoms, including positive (e.g., Hallucinogenic episodes) and negative (e.g., Anti-social tendencies). Finally, GWAS for AD cohorts has revealed multiple conserved genetic loci that could encode shared risk factors between the two diseases [10]. Clinical presentations, specifically psychiatric effects, create a crucial point of intersection to be explored where general psychosis was found in up to 60% of AD patients, including hallucination events as well as other effects mirroring those of the positive symptoms found in SCZ patients [11]. Thus, studying shared risk variants and genes between both diseases may help elucidate functional genomics of interest in both diseases. And it can further uncover cross-disease and disease-specific mechanisms between neurodegenerative and neuropsychiatric diseases.

Recently, advances in single-cell sequencing technologies have generated a great deal of excitement and interest in studying functional genomics at cellular resolution. For example, scRNA-seq and scATAC-seq techniques have measured the transcriptomics and epigenomics of individual cells in the human brain [12,13]. Further computational analyses have clustered cells into many cell types [12]. The cells in the same type share similar transcriptional activities such as gene expression and genomic functions. Differential gene expression across cell types is a complex, multi-gene dynamic process that tightly regulates and controls functions and is governed by gene regulatory factors such as transcriptional factors (TFs) and non-coding regulatory elements. These factors cooperate as a gene regulatory network (GRN) to facilitate the correct cellular and molecular functions on the genome-scale. Disrupted cooperation can give rise to abnormal gene expression, such as those present in diseases. Therefore, GRN has been used as robust system to infer genomic functions and molecular mechanisms, especially for human diseases [14].

Recent analyses have also revealed that brain disease risk variants are located in non-coding regulatory elements (e.g., enhancers). The risk genes likely have cell-type-specific effects for both neuronal and non-neuronal cell types [15,16]. Besides, recent single-cell studies suggest changes to cell-type-specific gene expression in brain diseases [8,9]. However, our understanding of the gene regulatory networks driving cell-type and disease-specific gene expression, especially across diseases, remains elusive. To address this issue, ones developed several computational methods to predict cell-type GRNs [17], such as PIDC [18], GENIE3 [19], and GRNBoost2 [20]. However, these methods typically only use single omics (e.g., transcriptomics) and predict networks based on statistical associations (e.g., co-expression), providing insufficient mechanistic insights into gene regulation at the cellular resolution. For instance, how the disease variants affect the transcription factor binding sites (TFBSs) on the distal regulatory elements (e.g., enhancers) that control disease genes is still unclear, especially at the cell-type level. Thus, it is essential to integrate emerging multi-omics data to understand cell-type gene regulation, especially involving non-coding regulatory elements. Recent studies have shown that integrating multi-omics data can reduce the impact of noise from a single omics data and achieve better prediction accuracy [21].

To explore these ideas, we developed a computational pipeline, scGRNom, by integrating multi-omics data to predict cell-type gene regulatory networks (GRNs) linking TFs, regulatory elements (e.g., enhancers and promoters), and target genes. In particular, we applied scGRNom to the multi-omics data at the cellular resolution, such as chromatin interactions, epigenomics, and single-cell transcriptomics of primary cell types in the human brain, including different excitatory and inhibitory neuronal types, microglia, and oligodendrocyte. Our predictions have high overlapping with state-of-the-art methods for revealing TFs and target genes [17], but they provide additional information on cell-type gene regulation, such as linking the regulatory elements to the genes. We also found that the enhancers in our cell-type GRNs are enriched with GWAS SNPs in human brain diseases, including psychiatric disorders and AD. Thus, we further linked the GWAS SNPs that interrupt TFBSs to cell-type disease genes based on the cell-type GRNs for SCZ and AD and found cross-disease and disease-specific genomic functions at the cell-type level. Finally, we found that the cell-type disease genes shared by AD and SCZ have improved predicting clinical phenotypes in AD, like disease staging and cognitive impairment.

## Methods and Materials

### Predicting gene regulatory networks from multi-omics data

scGRNom is a computational pipeline in R (https://github.com/daifengwanglab/scGRNom) to (I) integrate multi-omics datasets for predicting gene regulatory networks linking transcription factors, non-coding regulatory elements, and target genes, and (II) identify cell-type disease genes and regulatory elements. First, for predicting gene regulatory networks from multi-omics, scGRNom has three steps (**Figure 1**), each of which is available as an R function:

**Figure 1.**
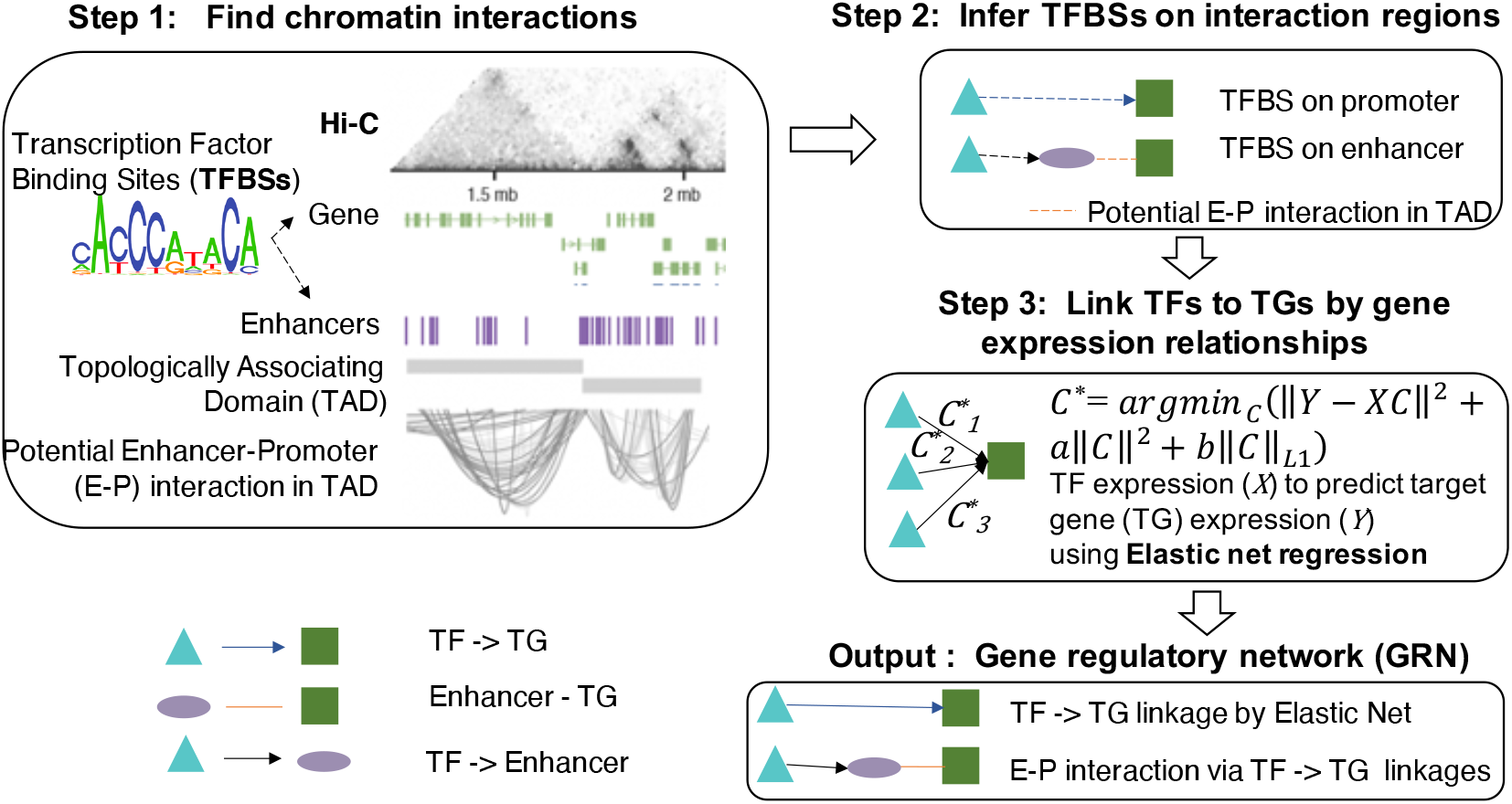
The computational pipeline, scGRNom, for predicting the gene regulatory network via multi-omics data. The pipeline inputs the chromatin interactions (e.g., from Hi-C) of regulatory elements (e.g., enhancer-promoter), identifies the transcription factor binding sites (TFBSs) on interacting regulatory elements, predicts TF-target gene expression relationships (e.g., high coefficients from Elastic net regression), and finally outputs a gene regulatory network linking TFs (cyan), regulatory elements (purple) to target genes (green).

#### Step1: Finding chromatin interactions

The function, *scGRNom_interaction,* inputs the chromatin interaction data (e.g., Hi-C) and predicts all possible interactions between enhancers and promoters in the data or the user-provided list - for example, those from Topologically Associating Domains (TADs) in Hi-C data. In addition, the function uses an R package, *GenomicInteractions* [22], to annotate interacting regions and link them to genes;

#### Step 2: Inferring the transcription factor binding sites on interacting regions

The function, *scGRNom_getTF,* infers the transcription factor binding sites (TFBS) based on consensus binding site sequences in the enhancers and promoters that potentially interact from the previous step *scGRNom_interaction*. It outputs a reference gene regulatory network linking these TF, enhancers, and/or promoters of genes. In particular, this function uses *TFBSTools* [23] to obtain the position weight matrices of the TFBS motifs from the JASPAR database [24] and predicts the TFBS locations on the enhancers and promoters via mapping TF motifs. The function further links TFs with binding sites on all possible interacting enhancers and promoters and outputs the reference regulatory network. Furthermore, this function can run on a parallel computing version via an R package, *motifmatchr* [25] for computational speed-up;

#### Step 3: Predicting the gene regulatory network

The function, *scGRNom_getNt* predicts the final gene regulatory network based on the TF-target gene expression relationships in the reference network. The reference gene regulatory network from the previous step provides all possible regulatory relationships (wires) between TF, enhancers, and target genes. However, the chromatin interacting regions are broad, so that many TFs likely have binding sites on them. Also, changes in gene expression may trigger different regulatory wires in the reference network. To refine our maps and determine the activity status of regulatory wires, we apply elastic net regression, a machine learning method that has successfully modeled TF-target gene expression relationships in the gene regulatory networks by our previous work [9]. Further, suppose the chromatin accessibility information is available (e.g., from scATAC-seq data for a cell type). In that case, the function can also filter the enhancers based on their chromatin accessibilities and then output the network links only having the enhancers with high accessibility (e.g., overlapped with scATAC-seq peaks). The parameter “open_chrom” inputs a list of user-defined chromatin accessible regions.

Mathematically, given a gene expression dataset and a reference network (e.g., from *scGRNom_getTF*), the function uses the TF expression to predict each target gene expression and finds the TF with high regression coefficients. Given a target gene, let 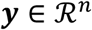 be a gene expression vector modeling its expression values across *n* samples (e.g., *n* cells from single cell data), and 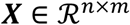 be the gene expression matrix of *m* TFs across *n* samples. Those *m* TFs should link to the target gene from the reference network, implying possible regulatory relationships to the gene. The elastic net regression model then aims to find the optimal coefficients for *m* TFs 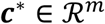 to solve the following optimization problem:

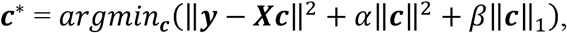

where *α* and *β* are parameters to adjust the contributions from L_2_ and L_1_ regularizations of 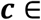 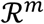. The samples are randomly divided into the training and testing sets by the parameter, train_ratio (e.g., if train_ratio=0.7, then 70% training and 30% testing data). The optimal TF coefficients, ***c***\* are estimated by the training data. Also, for the model evaluation, the mean square error (MSE) of the regression is calculated and reported by ∥***y***_***test***_ – ***X***_***test***_ ***c***\*∥^2^ using the testing data. Further, the top TFs with high coefficients can be either selected by absolute coefficient values (the parameter, cutoff_absolute) or a percentage from all *m* TFs (the parameter, cutoff_percentage). Finally, the function outputs a final gene regulatory network linking the top TFs as well as their linked enhancers (from the reference network, if any) to all possible target genes.

### Identifying cell-type disease genes and regulatory elements

In addition to predicting gene regulatory networks, the pipeline also provides another function, *scGRNom_disGenes,* for identifying cell-type disease genes and regulatory elements (e.g., enhancers, promoters). This function’s input includes a cell-type gene regulatory network and a list of GWAS SNPs associated with a disease (**Figure 2**). The function uses an R package, *GenomicRanges* [26], to overlap these disease SNPs with the enhancers and promoters of the input cell-type gene regulatory network, and then find the ones that interrupt the binding sites of all possible TFs (TFBSs) on the enhancers and promoters by *motifbreakR* [27]. It finally maps the overlapped enhancers or promoters and TFs with interrupted TFBSs onto the input network to find the linked genes and enhancers/promoters as the output cell-type disease genes and regulatory elements.

**Figure 2.**
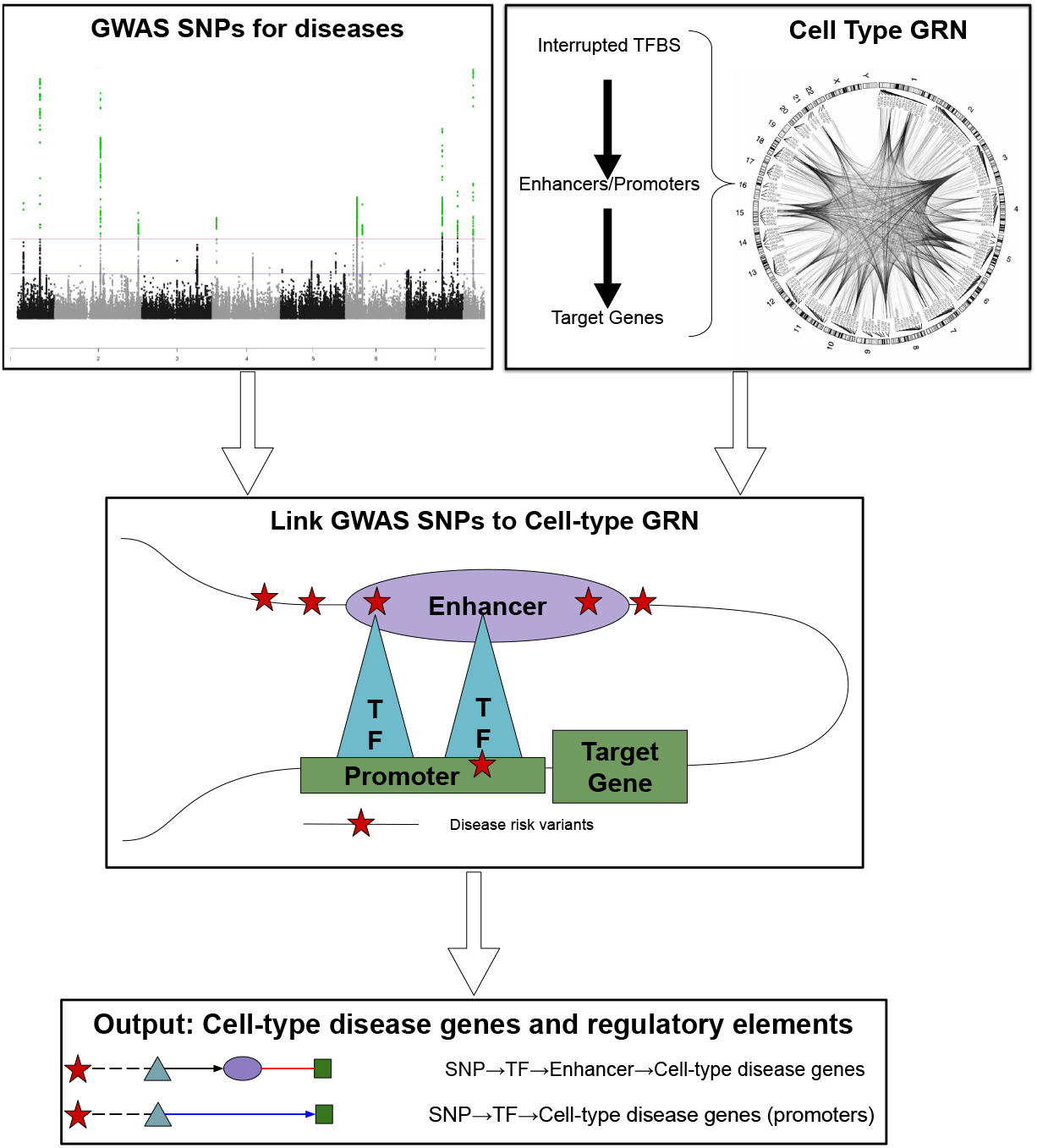
Identification of cell-type disease genes and regulatory elements. The function of our pipeline, *scGRNom_disGenes,* predicts cell-type disease genes and regulatory elements. First, it inputs a cell-type GRN (top right) and disease-associated SNPs from GWAS (top left). Second, the function identifies the disease SNPs that interrupt the binding sites of the TFs on the enhancers or promoters that link to the target genes in the input GRN (Middle). Finally, the function outputs a list of the target genes and regulatory elements (enhancers or promoters) linked by disease SNPs (Bottom). Red star: SNP. Cyan triangle: TF. Purple ellipse: Enhancer. Green square: cell-type disease gene.

### Application to multi-omics data and data processing for predicting cell-type gene regulatory networks in the human brain

We applied our computational pipeline to the multi-omics datasets at the cellular resolution in the human brain. We predicted cell-type gene regulatory networks for major cell types: excitatory and inhibitory neurons, microglia, and oligodendrocyte for the human brain [12]. The excitatory neuronal types include Ex1, Ex2, Ex3e, Ex4, Ex5b, Ex6a, Ex6b, Ex8, and Ex9. The inhibitory neuronal types include In1a, In1b, In1c, In3, In4a, In4b, In6a, In6b, In7, and In8. We first input recently published cell-type chromatin interactome data in the human brain [15] to *scGRNom_interaction* to reveal all possible interactions from enhancers to gene promoters in the neuronal, microglia, and oligodendrocyte types. The genome annotation was from TxDb.Hsapiens.UCSC.hg19.knownGene [28]. We then predicted a reference regulatory network for each of these cell types using *scGRNom_getTF*. Finally, given a cell type, we input the single cell gene expression data for the type and the reference network from *scGRNom_getTF* to *scGRNom_getNt* for predicting the cell-type gene regulatory network (GRN).

Specifically, to make our predicted networks comparable across cell types, we made the following data preparation and processing steps. First, although different studies have generated increasing numbers of single cell data (e.g., for microglia [8]), we used the data from one study (GSE97942) [12] that includes the gene expression data (UMI) of individual cells of primary cell types, all from one postmortem tissue of human frontal cortex, aiming not to introduce additional batches from different studies, which helps our further comparative analyses across cell types. In addition, we filtered the genes that express in less than 100 cells. We then normalized gene expression by *Seurat 4.0* [29] for further removing noises and batches across cell types. We also applied the method *MAGIC* [30] to impute the single cell gene expression of all cells to address potential dropout issues. Then for each cell type, we removed the lowly expressed genes with log10(sum of imputed gene expression levels of the cells of the cell type+1) < 1. The numbers of genes and cells for each cell type used for prediction are available in Supplemental File 1.

For predicting a cell type GRN using the Elastic net regression model, we randomly split the cells into 70% training and 30% testing sets. We then selected the best Elastic net model that minimized the mean square error (MSE). We then filtered the target genes based on the goodness of fit by MSEs. In particular, we removed the target genes predicted by Elastic net regression with MSE > 0.1 and also TF-target gene with absolute Elastic net coefficient < 0.01. The output cell-type GRN consists of the network edges that link TFs, enhancers (if any), and target genes (TGs), as well as the Elastic net coefficient of TF-TGs for each edge. Finally, we provided two versions of each cell-type GRN (Supplemental File 2):

I. the edges that include the enhancers that overlap the cell-type open chromatin regions predicted by recent scATAC-seq data (broad excitatory and inhibitory neurons, microglia and oligodendrocyte) [13];
II. the edges that only include the top 10% TFs with absolute coefficients for each target gene without considering cell-type open chromatin regions (for scATAC-seq data might be likely noisy and no open chromatin regions available for neuronal subtypes).

### Comparison with state-of-the-art methods

We compared our scGRNom predictions with existing state-of-the-art methods. In particular, we input the single-cell gene expression data for each cell type to a recently published benchmark framework, BEELINE [17], and predicted the cell-type regulatory networks using three of the most consistent and highly accurate methods, PIDC, GENIE3, and GRNBoost2. These methods only input gene expression data to predict all possible TF-target gene (TG) regulatory links based on their expression relationships, without considering the regulatory elements or open chromatin regions at the cell-type level. Thus, after applying to our processed single cell gene expression data, they generated more network edges than us because scGRNom only keeps the TF-TG links in which TFs have binding sites on the regulatory elements (e.g., enhancers and promoters). Therefore, to make these networks comparable with ours, we extended our networks by selecting up to the top 30% TFs for each TG and then checked if our TF-TG links predicted by the state-of-the-art method (also up to top 30% TFs for each TG selected for each method). In particular, given a cell-type GRN by scGRNom, we selected top K TFs per target gene (TG) to see if the TF-TG pairs were also predicted by PIDC, GENIE3, or GRNBoost2 (again picking top K TFs per TG). We varied K values from 0% to 30% and then calculated the percentages of the scGRNom’s TF-TG pairs that can be predicted by one of those methods (Figure S1).

### GWAS SNPs and heritability enrichment analyses for the enhancers in the cell-type gene regulatory networks in the human brain

Genome Wide Association Studies (GWAS) have identified a variety of genetic risk variants, including single nucleotide polymorphisms (SNPs) that are significantly associated with diseases and phenotypes (i.e., the traits). For example, recent GWAS studies have identified many SNPs associated with Alzheimer’s Disease (AD, 2357 credible SNPs) [6] and Schizophrenia (SCZ, 6105 credible SNPs) [7]. In addition to the credible SNPs, we also included additional SNPs with *p*<5e-5 from the AD and SCZ GWAS summary statistics for linking potential additional cell-type disease genes [31]. We applied the *partitioned linkage disequilibrium score regression (LDSC)* [32] to evaluate the heritability explained by the enhancers of our cell-type gene regulatory networks for GWAS SNPs. In particular, our heritability enrichment analyses used the GWAS summary statistics for the diseases or traits: Schizophrenia (SCZ) [7], Alzheimer’s disease (AD) [6], Autism spectrum disorder (ASD) [33], Bipolar disorder (BPD) [34], Amyotrophic lateral sclerosis (ALS) [35], Major depressive disorder (MDD) [36], Intelligence [37], Multiple sclerosis (MS) [38], Parkinson’s disease (PD) [39], Attention deficit hyperactivity disorder (ADHD) [40], Education [41], Type 2 diabetes (T2D) [42], Inflammatory bowel disease (IBD) [43], Coronary artery disease (CAD) [44]. Also, we provided the numbers of GWAS SNPs for AD and SCZ that interrupt the binding sites of at least one of all possible TFBSs and the binding sites of the regulatory TFs in each cell-type GRN (Table S1).

### Identification and enrichment analyses of cell-type disease genes

Given a cell-type and a disease type (AD or SCZ), we input the cell-type GRNs (both versions) and the GWAS SNPs for the disease to the *scGRNom_disGenes* function for identifying the cell-type disease genes. We identified the cell type disease genes (AD and SCZ) for all the cell types as above, including excitatory and inhibitory neuronal subtypes, microglia, and oligodendrocyte (Supplemental File 3). We also merged the disease genes for excitatory/inhibitory neuronal subtypes as the broad excitatory/inhibitory neuronal disease genes. We used the web app, *Metascape* [45] to find the enrichments of cell-type disease genes such as KEGG pathways, GO terms, protein-protein interactions, and diseases (via DisGeNET). Enrichment p-values shown in this paper were adjusted using the Benjamin-Hochberg (B-H) correction. Also, we looked at the expression levels of our cell-type disease genes in the disease samples. In particular, we compared the published population gene expression data in AD for single cell expression [8].

### Machine learning prediction of clinical phenotypes from cell-type disease genes

Finally, we used the machine learning approach to predict clinical phenotypes from our cell-type disease genes using the population data of the ROSMAP project, an independent AD cohort [46]. Given a clinical phenotype, we assume that *X*_*i*_ ∈ *R*^*d*^ represents the expression data of *d* disease genes for the *i*^th^ individual in the cohort and *Y*_*i*_ ∈ {0,1} represents the binarized class of the *i*^th^ individual’s phenotype with *i* ∈ 1,…, *n* individuals for training. We then found the optimal logistic regression model with the parameters 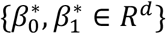 to classify the phenotype from the disease gene expression data via minimizing the following loss function:

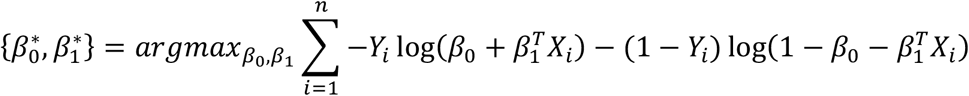

where (.)^*T*^ is the transpose operation. Also, we performed Cross-Validation (*K*=5) for the individual samples with 80% training and 20% testing sets. We also balanced the class sample size in each training set by the weighting method [47] so that the baseline of the classification accuracy is 50% for two classes. We used the individuals from the training sets to train the classification model. We then used the individuals from the testing sets to evaluate the classification performance, i.e., accuracy. In particular, we predicted four clinical phenotypes in ROSMAP: Braak Stages that measure the severity of neurofibrillary tangle (NFT) pathology (Braak early stages (0,1,2,3) vs. late stages (4, 5, 6)), CERAD scores that measure neuritic plaques (no AD vs. AD), the diagnosis of cognitive status (DCFDX, no or mild cognitive impairments (1,2,3) vs. Alzheimer’s dementia (4, 5)) and the cognitive status at the time of death (COGDX, no or mild cognitive impairments (1,2,3) vs. Alzheimer’s dementia (4, 5)).

## Results

### Predicting cell-type gene regulatory networks in the human brain

We applied the scGRNom pipeline to the multi-omics data for the human brain, including cell-type chromatin interactions [15], transcription factor binding sites [23], single-cell transcriptomics [9] and recent cell-type open chromatin regions [13] (Methods and Materials). In particular, we predicted the cell-type gene regulatory networks (GRNs) for both glial and neuronal types, including microglia, oligodendrocyte, excitatory neuronal subtypes (Ex1, Ex2, Ex3e, Ex4, Ex5b, Ex6a, Ex6b, Ex8 and Ex9, and inhibitory neuronal subtypes (In1a, In1b, In1c, In3, In4a, In4b, In6a, In6b, In7, and In8). Each cell-type GRN links TFs to enhancers to target genes (TGs), and has two versions that (I) only include the network edges with open cell-type enhancers predicted by recent scATAC-seq data [13] and (II) only include the edges with top 10% TFs with highest absolute coefficients for each target gene without using cell-type open chromatin regions (due to scATAC-seq data might be likely noisy and not specific to neuronal subtypes). For instance, we found that the microglia GRN with open chromatin regions consists of 47353 edges linking 180 TFs, 1893 microglia open enhancers, 1236 TGs. All cell-type GRNs are available in Supplemental File 2. The network statistics such as numbers of cells, edges, TFs, enhancers and TGs for all cell types are available in Supplemental File 1.

Our GRNs reveal many known cell-type-specific regulations. **Figure 3A** visualizes the subnetworks among TFs for select cell types (i.e., TGs are also TFs). For example, two known TFs, MEF2A and RFX3, that control microglia phenotypes play hub roles in the microglia network [48,49]. The nuclear factors NFIA, NFIX, and FOXP1 controlling neural differentiation and gliogenesis are also hub genes in the oligodendrocyte network [50–52]. MEF2C regulating inhibitory and excitatory synapses is a central node in both excitatory and inhibitory networks (e.g., Ex1 and In6b) [53]. In addition to cell-type TFs, we also observed the cell-type-specific expression relationships between TFs and TGs (high correlation). For instance, in **Figure 3B**, E2F3-LRRK2, STAT2-FBXO32, IRF2-DYNC1, and ATF4-EPB41L1 show cell-type-specific expression relationships across the cells of microglia, oligodendrocyte, Ex1, and In6b types.

**Figure 3.**
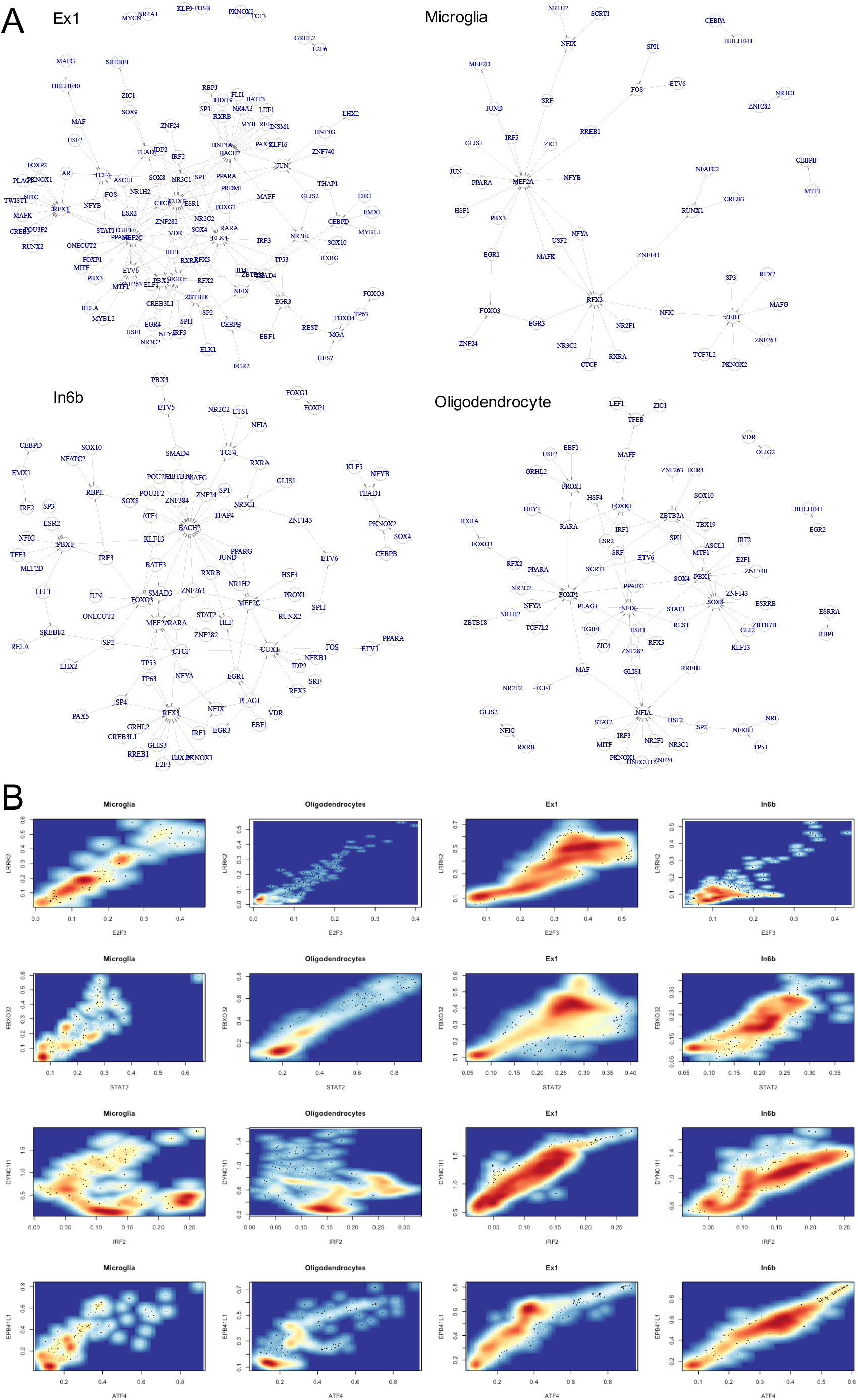
Cell-type gene regulatory networks (GRNs) in the human brain. (**A**) The subnetworks of select cell-type GRNs among TFs (i.e., TG is also TF): Ex1 (Top left), Microglia (Top right), In6b (Bottom left), Oligodendrocyte (Bottom right). (**B**) The expression levels of the cells for select TFs and TGs across the four cell types in (**A**). TF: x-axis. TG: y-axis. The select TF-TG pairs from top to bottom are E2F3-LRRK2, STAT2-FBXO32, IRF2-DYNC1, and ATF4-EPB41L1. The red darkness corresponds to the expression level.

In addition, we compared our predicted cell-type gene regulatory networks with existing state-of-the-art methods for predicting cell-type gene regulatory networks, particularly those that are consistent and highly accurate PIDC, GENIE3, and GRNBoost2 benchmarked by BEELINE [17] (Materials and Methods). The percentages of the overlapped TF-TG links of the cell-type network between scGRNom (both versions) and the state-of-the-art methods are over 50% (Methods, **Figure S1**). This suggests a high consistency between scGRNom and these methods. However, these methods predicted TF-TG links without providing information on regulatory elements like enhancers. Thus, we looked further at the enhancers in our cell-type GRNs and found that they have significantly high heritability enrichments for GWAS SNPs of multiple brain diseases and traits (*p*<0.05, Materials and Methods). For example, the enhancers in our excitatory-neuron and inhibitory-neuron GRNs have high enrichments for AD, SCZ, Major depressive disorder, intelligence, education (**Figure 4A, Figure S2**). Besides, these brain-cell-type enhancers do not have significant enrichment of GWAS SNPs for non-brain diseases. Therefore, the heritability enrichment analysis suggests that the enhancers in our cell-type GRNs have potential pleiotropic roles associated with multiple brain diseases or traits. Finally, we also compared our cell-type networks with public GRN databases such as TRRUST [54], Dorothea [55], and RegNetwork [56]. Those GRNs were primarily inferred by integrating different studies from the literature (e.g., via physical interactions, co-expressed genes at the bulk tissue level) and thus may not be specific for the neuronal and glial cell types in the human brain. However, we still found many overlapped network edges (TF-TG) with our cell-type gene regulatory networks. For example, we found that 319 TRRUST edges, 5935 Dorothea edges, 106 RegNetwork edges overlap with at least one of our cell-type GRNs, implying their potential associations with the human brain’s cell types.

**Figure 4.**
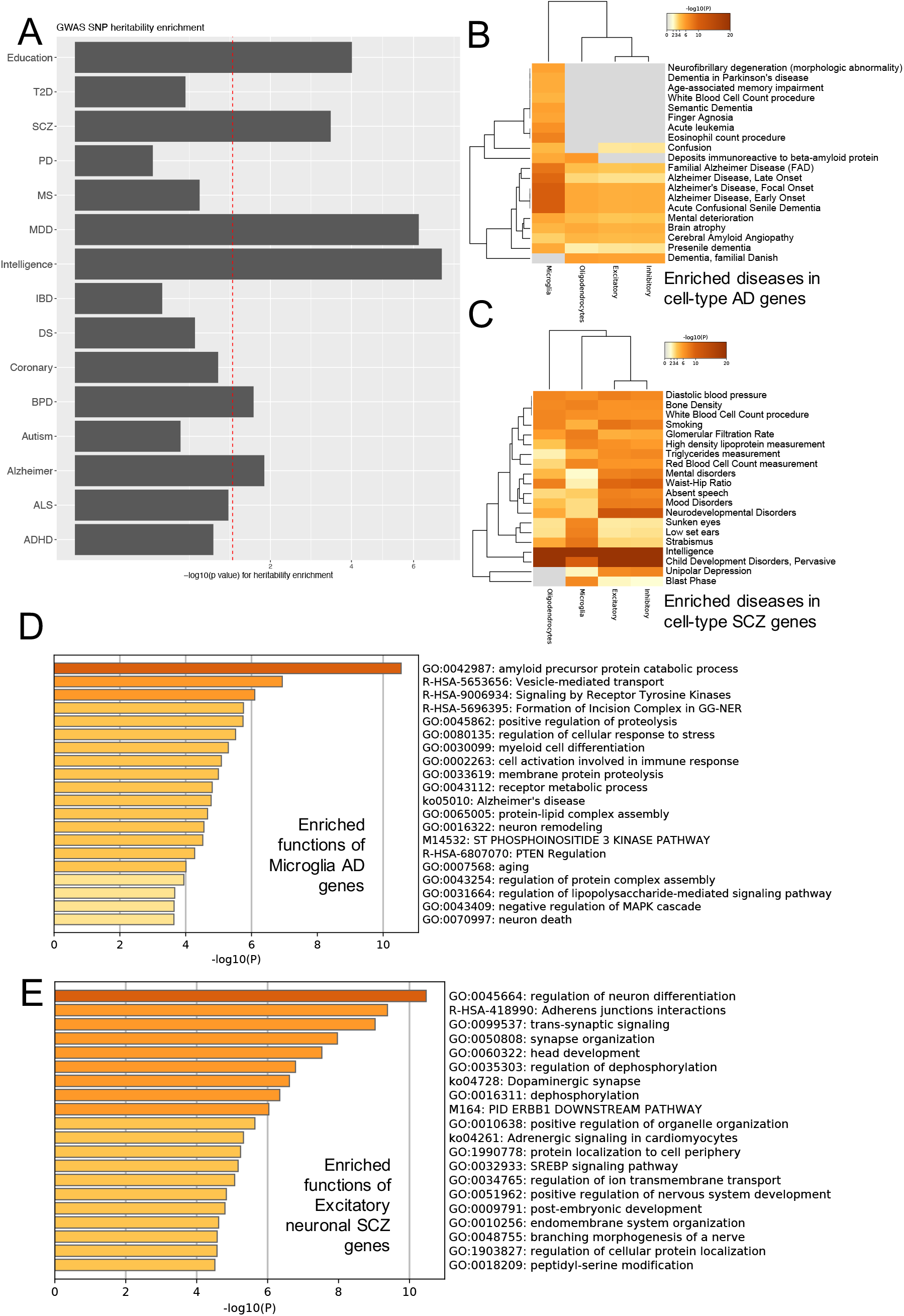
Enrichments of cell-type enhancers and disease genes. (**A**) Partitioned heritability enrichment of GWAS SNPs associated with various diseases and traits (bar) on the enhancers of In1a GRN. Bar height is −log10(*p* value) of the enrichment. The diseases and traits are Schizophrenia (SCZ), Alzheimer’s disease (AD), Autism spectrum disorder (ASD), Bipolar disorder (BPD), Amyotrophic lateral sclerosis (ALS), Major depressive disorder (MDD), Intelligence, Multiple sclerosis (MS), Parkinson’s disease (PD), Attention deficit hyperactivity disorder (ADHD), Education, Type 2 diabetes (T2D), Inflammatory bowel disease (IBD), Coronary artery disease (CAD). The red line represents *p* value = 0.05. (**B**) The disease enrichments of cell-type AD genes. Rows are diseases/traits. Columns are cell types. Darkness is proportional to −log10(p) of the enriched term. (**C**) The disease enrichments of cell-type SCZ genes. Rows are diseases/traits. Columns are cell types. Darkness is proportional to −log10(p) of the enriched term. (**D**) The enrichments of functions and pathways (e.g., GO, KEGG, REACOME) in the microglia AD genes. Bar darkness and length are proportional to −log10(p) of the enriched term. (**E**) The enrichments of functions and pathways (e.g., GO, KEGG, REACOME) in the excitatory neuronal SCZ genes. Bar darkness and length are proportional to −log10(p) of the enriched term.

### Identifying cell-type disease genes in AD and SCZ for neuronal and glial types

We used these cell-type GRNs to link GWAS SNPs with disease risk genes for each cell type, advancing knowledge on cross-disease and disease-specific interplays among genetic, transcriptional, and epigenetic risks at cellular resolution. In particular, we chose SCZ and AD, two majorly represented neuropsychiatric and neurodegenerative diseases with potential convergent underlying mechanisms [4], and we linked a number of cell-type disease genes (Methods, Supplemental File 3) and performed their enrichments (Supplemental File 4). We found that many disease genes present in one or a few cell types only, suggesting potential cell-type-specific contributions to AD and SCZ (**Figure S3**). As shown in **Figure 4B**, the cell-type AD genes are also significantly enriched with known disease genes for AD and other dementia diseases such as brain atrophy and cerebral amyloid angiopathy (p<0.01). For example, the microglia AD genes, including APP, CLU, BACE2, and BIN1, are specifically enriched for other diseases such as neurofibrillary degeneration, Parkinson’s dementia, and aging memory impairment. Also, the cell-type SCZ genes are enriched for various disorders such as neurodevelopmental, mental and mood disorders, and depression (*p* < 0.01, **Figure 4C**). Furthermore, many cell-type disease genes have corresponding expression activities in the disease samples. For example, 72 excitatory AD genes (86%) and 65 inhibitory AD genes (80%) (plus 22 oligodendrocyte AD genes and three microglia AD genes) are significantly differentially expressed in the corresponding cell types in AD individuals, respectively (*p* < 0.05) [8].

The functional enrichment analyses for our cell-type disease genes also uncover known genomic functions and pathways at the cell-type level. For instance, the microglia AD genes are enriched with amyloid beta formation and clearance [57], MAPK signaling [58] and neuron death [59] (**Figure 4D**), and the oligodendrocyte AD genes are enriched with Tau protein binding [60]. This is vital to understanding a multitude of diseases that commonly demonstrate atrophy of cortical tissue as a hallmark feature. In SCZ genes, we also found that multiple key hallmark pathways were enriched, such as dopaminergic synapse [61], trans-synaptic signaling [62], and synapse organization [63] for excitatory SCZ genes (**Figure 4E**). For inhibitory SCZ genes, we observed that MAPK family signaling [64], regulation of NMDA receptor activity [65], dopaminergic synapses [61] and neurotransmission [65] are enriched.

### Comparative analyses reveal the interplays between genomic functions, pathways, cell types, and diseases

In addition to cell-type-specific pathways in these diseases, we also identified those involving multiple cell types in each disease, implying that potential cell-type interactions drive the disease pathology. For example, the enrichment of SCZ primarily includes changes in synapse structures and cell shaping and differentiation (**Figure 5A**). Clinically, this is consistent with the consensus that SCZ is strictly neuropsychiatric as opposed to degenerative. In particular, cell morphogenesis and regulation of neuron differentiation are enriched in all four major cell type SCZ genes (*p* < 0.01). Early life neurodevelopmental genetic markers may suggest causal links with alterations in hippocampal cell differentiation points on the front of cell morphogenesis, leading to cascades of downstream effects [66]. This has primarily been studied and modeled within the scope of iPSC-based analyses, which make correlations and connections to the clinical presentation more difficult due to the additional abstraction from the standard pathology-based analysis. Also, the BDNF signaling pathway that potentially relates to intercellular communications is enriched as well in multiple cell types (*p* < 0.01) [67]. Finally, we also observed that protein-protein interactions (PPIs) are enriched among the disease genes of the cell types at a higher level. As shown in **Figure 5B**, the SCZ genes for dopaminergic synapse, NMDA receptors, glutamate binding, and activation are shared by multiple cell types and have strong PPIs, implying protein-level cross-type coordination [68]. In AD, multiple pathways were significantly enriched across various cell types (**Figure 5C**). For instance, the catabolic process for the AD key player, amyloid precursor protein (APP) is enriched with both glial and neuronal types (*p* < 0.01) [69].

**Figure 5.**
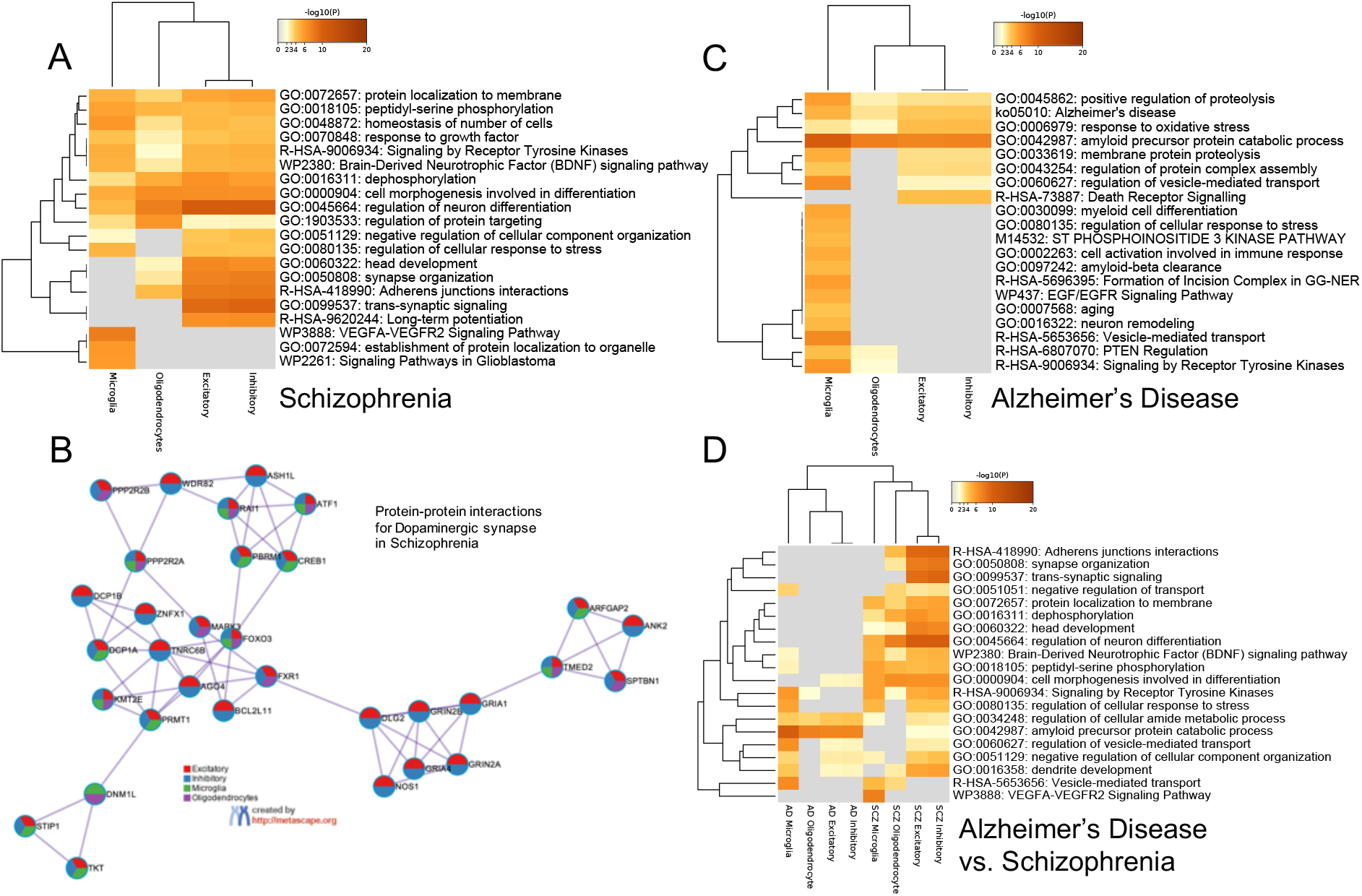
Cross-cell-type conserved and cell-type-specific functions, pathways and protein-protein interactions in Schizophrenia and Alzheimer’s Disease. Darkness in heatmaps is proportional to −log10(p) of the enrichment. (**A**) The enrichments of select conserved and specific functions/pathways (e.g., GO, KEGG, REACTOME) across cell-type disease genes in Schizophrenia. (**B**) The enrichments of protein-protein interactions among major cell-type disease genes in Schizophrenia: Excitatory neuron (broad, red), Inhibitory neuron (broad, blue), Microglia (green), Oligodendrocyte (Purple). (**C**) The enrichments of select conserved and specific functions/pathways across cell-type disease genes in Alzheimer’s Disease. (**D**) The enrichments of select conserved and specific functions/pathways across cell-type disease genes between Schizophrenia and Alzheimer’s Disease.

Furthermore, we found that cross-disease conserved functions/pathways are involved in one or multiple cell types, revealing potential novel functional interplays across cell types and diseases (**Figure 5D**). For example, the cell morphogenesis involved in differentiation is enriched in both AD and SCZ neuronal genes. Another example is that the vesicle-mediated transport is enriched for both AD and SCZ microglia genes. In total, we found 11 microglia genes shared by AD and SCZ. In particular, the VEGF signaling pathway is enriched in the SCZ microglia genes. In the general theme of AD pathology, increased VEGF expression has been linked to worse cognitive outcomes in post-mortem analysis [70]. Similarly, multiple meta analyses have revealed differential expression levels between healthy controls and SCZ patients [71]. However, little has been done to link potential gene function to cell-type level interactions and pathways. Here, these SCZ-AD shared microglia genes may help explain shared higher-level dysfunction between both diseases as evidenced by higher expression. Also, we found that some functions involve different cell types across diseases. The dendrite development has been found in both SCZ and AD pathology [72,73]. We found that it is mainly enriched with microglia in AD but neuronal types in SCZ.

More interestingly, when exploring the interactions between cell types that change between diseases, the disease-specific pathologies enter to explain the cause of discrepancies. In particular, for AD, it is shown that phagocytic microglia are activated during the early stages of synaptic decline, leading to eventual neuroinflammation and programmed cell death [74]. For SCZ, the oligodendrocyte enrichment reveals similar intercellular mechanisms between excitatory and inhibitory neurons, specifically those regulating neuron differentiation (**Figure 5A**), providing potential direction for future exploration and validation of the communication role of oligodendrocytes [75].

### Prediction of clinical phenotypes from cell-type disease genes

Finally, we want to investigate the clinical applications of our cell-type disease genes. To this end, we looked at the population-level gene expression data for Alzheimer’s disease in the ROSMAP cohort [46]. In particular, we first found that many cell-type AD genes have significantly associated expression levels with clinical phenotypes across individuals in ROSMAP. For example, out of 195 cell-type AD genes, we found that 72 genes are significantly associated with the Braak Stages that measure the severity of neurofibrillary tangle (NFT) pathology (ANOVA *p* < 0.05), 78 genes with the CERAD scores that measure neuritic plaques (ANOVA *p* < 0.05), 89 genes with the diagnosis of cognitive status (DCFDX) (ANOVA *p* < 0.05), 73 genes with the cognitive status at the time of death (COGDX) (ANOVA *p* < 0.05) and 92 genes with the Mini-Mental State Examination (MMSE) scores (Pearson correlation *p*< 0.05). In total, 135 cell-type AD genes were significantly associated with at least one clinical phenotype in ROSMAP (Supplemental File 5).

In addition to statistically significant associations between our cell-type disease genes and clinical phenotypes, we also applied the machine learning approach to predict clinical phenotypes from these cell-type disease genes using ROSMAP data (Methods). Specifically, for each clinical phenotype, we used the logistic regression model to classify individual states of the phenotype (as classes) from their expression data of 53 AD-SCZ shared cell-type genes. We performed the cross-validation (*K*=5) for the individual samples with 80% training and 20% testing sets. As shown in Fig. 6, our average accuracy values for classifying all four major clinical phenotypes: Braak Stages, CERAD scores, DCFDX status, COGDX status are much higher than the baselines (50%), the random select genes, and AD genes from the latest GWAS study [6]. This suggests that using cell-type disease genes shared by AD and SCZ has improved predicting those clinical phenotypes, especially for cognitive related ones in AD.

**Figure 6.**
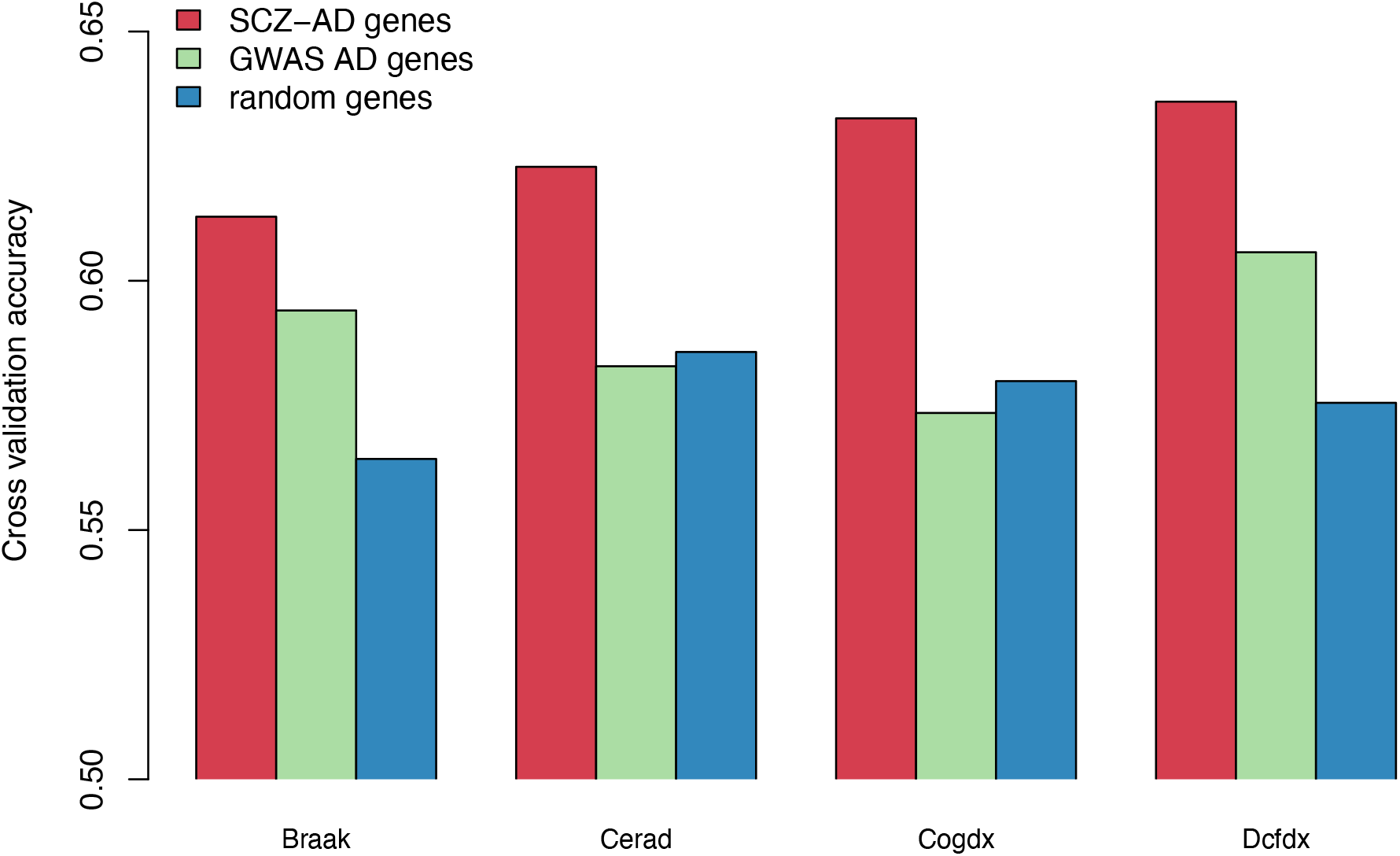
Prediction accuracy of AD clinical phenotypes from disease genes. The AD population data for prediction was from the ROSMAP cohort [46]. AD clinical phenotypes include Braak – stages that measure the severity of neurofibrillary tangle (NFT) pathology, Cerad – scores that measure neuritic plaques, Cogdx – cognitive status at the time of death, and Dcfdx - the diagnosis of cognitive status. The bar height represents the average accuracy of cross-validation (*K*=5) from the prediction using logistic regression (Methods). Red: scGRNom’s cell-type disease genes shared by AD and SCZ (SCZ-AD genes). Green: AD genes from GWAS [6]. Blue: randomly select genes (same number as SCZ-AD genes).

## Discussion

This paper focuses on the scGRNom’s application to single cell data for Alzheimer’s and schizophrenia. However, scGRNom is general-purpose for understanding functional genomics and gene regulation in other diseases. Besides the cell types, the pipeline also predicts the gene regulatory networks for cell clusters (unknown cell types) and bulk tissue types. Furthermore, recent eQTL studies have identified various SNPs associating with gene expression in multiple brain tissue types using the population data such as GTEx [76] and PsychENCODE [9]. Although those brain eQTLs suggest the SNP-gene association at the bulk tissue level, we still found that several eQTLs in PsychENCODE match our linked GWAS SNPs and cell-type disease genes, e.g., SNPs chr2:127846321 for and chr20:43598154 for STK4, two microglia disease genes for AD and SCZ, respectively. This suggests potential cell-type effects of these human brain eQTLs. Thus, increasing single cell data at the population level allows us to predict the cell-type eQTLs [77], which will likely help understand cell-type gene regulation and refine linking disease genes at the cell-type level.

For linking disease genes, we primarily used the interrupted TFBSs by GWAS SNPs. Future studies utilizing scGRNom would be able to take advantage of the ever-growing number of GWAS for a wide variety of diseases as well as single cell data. However, additional information can also help link GWAS SNPs to disease genes. For example, existing tools such as FUMA [78] have linked GWAS loci to genes by integrating information from multiple resources, providing more functional linkages from genotype to genes to phenotype in human diseases. Thus, incorporating multiple GWAS data (e.g., various brain regions) exposes key areas of observed phenotypes. Previous studies have demonstrated the caution that must be exercised when attempting to correlate GWAS data with clinical phenotypes, and methods such as our analysis mitigate these effects [16]. A similar methodology as outlined could be used where common loci within each set of summary statistics are incorporated and established before integration into the cell type GRNs, thus linking neuronal spatial information with known mutation sites in patients along with potentially cell-type-specific functionality. Expanding past the cell types examined here into additional multicellular analyses is also possible given expanded interactome data. Notably, this would allow for further investigation of complex neuropsychological diseases and cases where the line between different clinical classifications becomes blurred and leads to additional complications about clinically relevant genetic therapies. One such example includes Autism Spectrum Disorder (ASD), where clinical presentations can vary in multiple axes of severity, creating a broad spectrum of phenotypes. In such cases, potentially linking specific symptoms or aspects of a particular subset of ASD to particular brain regions and cell types allows for a better-informed picture of functional consequences associated with genetic mutation sites. Such connections could aid in determining genetic risk factors associated with variations in edge case patients; they also create the opportunity to take advantage of Induced Pluripotent Stem Cell (iPSC) technology using genetic engineering technologies to create point mutations matching computationally identified genes. Moreover, because the single cell multi-omics data we used for predicting cell-type GRNs are not specific for particular diseases (AD or SCZ), we used the SNPs disrupting all possible TFs on the enhancers and promoters from our cell type GRNs to link at large cell-type disease genes. However, the scGRNom pipeline is general-purpose and able to work for incoming disease-specific single cell multi-omics, and link to cell-type disease genes via interrupted regulatory TFs in the diseases.

Machine learning has also been widely used to analyze multi-omics, such as multiview learning and deep learning [9,79]. Multiview learning has great potential for understanding functional multi-omics and revealing nonlinear interactions across omics. Therefore, integrating such emerging machine learning approaches will enable identifying different cross-omic patterns, especially for increasing single-cell multi-omics data and providing more comprehensive mechanistic insights in cell-type gene regulation and linking to disease genes. For example, this means adding more omics such as methylation data that reflect epigenetic changes that may occur due to wide variations of inherited and environmental factors [80]. At a deeper functional level, variations in methylation have been attributed to alterations in splicing activity, ultimately impacting the regulation and expression of key genes [81]. Additionally, integrating proteomic data at a single-cell level enhances the broader picture formed through additional data sources even further [82]. Lastly, expanding past simple methylation and proteomics allows for the ability to include all forms of data incorporated through the single-cell cytometry [83].

## Conclusions

We developed a computational pipeline, scGRNom, to integrate multi-omics data and predict gene regulatory networks (GRNs), which link TFs, non-coding regulatory elements (e.g., enhancers), and target genes. With applications to the data from single-cell multi-omics of the human brain, we predicted cell-type GRNs for both neuronal (e.g., excitatory, inhibitory) and glial cell types (e.g., microglia, oligodendrocyte). Further, scGRNom can input cell-type GRNs and disease risk variants to link disease genes at the cell-type level, such as brain diseases like Alzheimer’s and schizophrenia. These disease genes revealed conserved and specific genomic functions across neuropsychiatric and neurodegenerative diseases, providing cross-disease regulatory mechanistic insights at the cellular resolution. Although this paper focuses on Alzheimer’s and schizophrenia, scGRNom is general-purpose for understanding functional genomics and gene regulation in other diseases, at either bulk tissue or cell-type levels. Finally, scGRNom is open-source available at https://github.com/daifengwanglab/scGRNom.

## Supplementary information

Supplemental materials – supplemental figures and table

Supplemental File 1 – statistics of cell-type gene regulatory networks

Supplemental File 2 – cell-type gene regulatory networks (TF, enhancer, target gene) Supplemental File 3 – cell-type disease genes for AD and SCZ

Supplemental File 4 – enrichments of cell-type disease genes for AD and SCZ

Supplemental File 5 – cell-type disease genes associated with clinical phenotypes in AD

## Data availability

Our pipeline for predicting gene regulatory networks via multi-omics is open-source available, as well as a tutorial at https://github.com/daifengwanglab/scGRNom. All data supporting this study are included in this paper, the supplemental files, and/or available from the corresponding author on request.

## Author contributions

D.W. conceived and designed the study. T.J. and M.Y. implemented the software. T.J., P.R., M.Y., J.H., S.L., and D.W. analyzed the data. T.J., P.R., M.Y., Pa.R., and D.W wrote the manuscript. All authors read and approved the final manuscript.

